# Characterisation of the biosurfactants from phyllosphere colonising *Pseudomonads* and their effect on plant colonisation and diesel degradation

**DOI:** 10.1101/2020.10.27.358416

**Authors:** S Oso, F Fuchs, C Übermuth, L Zander, S Daunaraviciute, DM Remus, I Stötzel, M Wüst, L Schreiber, MNP Remus-Emsermann

**Affiliations:** School of Biological Sciences, University of Canterbury, Christchurch, New Zealand; Department of Institute for Cellular and Molecular Botany (IZMB), University of Bonn, Bonn, Germany; Protein Science & Engineering, Callaghan Innovation, School of Biological Sciences, University of Canterbury, Christchurch, New Zealand; Institute of Nutritional and Food Sciences (IEL), University of Bonn, Bonn, Germany; Biomolecular Interaction Centre, Christchurch, New Zealand; Bio-Protection Research Centre, School of Biological Sciences, University of Canterbury, Christchurch, New Zealand

**Keywords:** Plant-microbe interactions, phyllosphere-inhabiting microbes, leaf wax, leaf cuticle, aliphatics, viscosin, Massetolide A

## Abstract

Biosurfactant production is a common trait in leaf surface colonising bacteria that has been associated with increased survival and movement on leaves. At the same time the ability to degrade aliphatics is common in biosurfactant-producing leaf colonisers. Pseudomonads are common leaf colonisers and have been recognised for their ability to produce biosurfactants and degrade aliphatic compounds. In this study, we have investigated the role of biosurfactants in four non-plant plant pathogenic *Pseudomonas* strains by performing a series of experiments to characterise the surfactant properties, and their role during leaf colonisation and diesel degradation. The produced biosurfactants were identified using mass-spectrometry. Two strains produced viscosin-like biosurfactants and the other two produced Massetolide A-like biosurfactants which aligned with the phylogenetic relatedness between the strains. To further investigate the role of surfactant production, random Tn*5* transposon mutagenesis was performed to generate knockout mutants. The knockout mutants were compared to their respective wildtypes in their ability to colonise gnotobiotic *Arabidopsis thaliana* and to degrade diesel. It was not possible to detect negative effects during plant colonisation in direct competition or individual colonisation experiments. When grown on diesel, knockout mutants grew significantly slower compared to their respective wildtypes. By adding isolated wildtype biosurfactants it was possible to complement the growth of the knockout mutants.

**Importance:** Many leaf colonising bacteria produce surfactants and are able to degrade aliphatic compounds, however, if surfactant production provides a competitive advantage during leaf colonisation is unclear. Furthermore, it is unclear if leaf colonisers take advantage of the aliphatic compounds that constitute the leaf cuticle and cuticular waxes. Here we test the effect of surfactant production on leaf colonisation and demonstrate that the lack of surfactant production decreases the ability to degrade aliphatic compounds. This indicates that leaf surface dwelling, surfactant producing bacteria contribute to degradation of environmental hydrocarbons and may be able to utilise leaf surface waxes. This has implications for plant-microbe interactions and future studies.

## Introduction

The leaf cuticle is a hydrophobic barrier which consist of cutin, a polymer of very long chain aliphatics, interspersed and overlaid by very long chain monomeric aliphatics, cuticular waxes (Kolattukudy, 1980; Zeisler-Diehl et al., 2018). The cuticle reduces water loss, provides protection against UV radiation, and is the primary interface for plant microorganism and insect interactions (Riederer & Schreiber, 2001; Serrano et al., 2014; Yeats et al., 2012). The cutin is a biopolymer which consists mainly of ω −and midchainhydroxy and epoxy fatty acids C_16_-C_18_ as well as glycerol (Graça, 2002; Pollard et al., 2008; Wattendorff & Holloway, 1980). The cutin forms the structural backbone of the cuticle as it is known to prevent mechanical damage. The cuticular waxes are the second major component of the leaf cuticle mostly consisting of alkanes, alcohols, acids, and aldehydes of chain lengths between C_16_ - C_32_ Cuticular waxes may also include secondary metabolites such as flavonoids, triterpenoids and phenylpropanoids (Jeffree, 2006). Cuticular waxes can be separated into two distinct waxes. The intracuticular wax within the cutin polymer is clearly distinct from the epicuticular wax which is on the outer surface of the cutin polymer (Buschhaus & Jetter, 2011; Samuels et al., 2008). These differences thus affect the physical properties of the plant surfaces. The composition of the cuticular waxes is dependent on plant species and environmental conditions (Jetter et al., 2006; Shepherd & Wynne Griffiths, 2006). Wax monomers are very energy rich and a potential source of energy and carbon if they are bioavailable. However, it is still unclear if bacteria are able to utilise these aliphatic compounds constituting the cuticle of living leaves as a source of carbon and if surfactants would facilitate the utilisation.

Leaves are home to a manifold of bacteria and they can be covered by up to 5% bacterial biomass (Remus-Emsermann et al., 2014; Schlechter et al., 2019). Many leaf surface colonising genera were previously shown to degrade hydrocarbons, e.g. *Rhodococcus* spp., *Sphingomonas* spp., *Pantoea* spp., *Methylobacterium* spp., and Pseudomonads (Kertesz & Kawasaki, 2010; Oso et al., 2019; Pizzolante et al., 2018; Salam et al., 2015). Pseudomonads are common leaf colonisers and have many different ecological roles, e.g. many *Pseudomonas syringae* strains can be bonafide and host specific pathogens (Xin et al., 2018) while others may act as antagonists against agents of plant disease (Cabrefiga et al., 2007; Zengerer et al., 2018) or have unknown, tritagonistic (Freimoser et al., 2016), functions in the microbiota (Remus-Emsermann et al., 2016; Schmid et al., 2018). Pseudomonads have the ability to produce so-called biosurfactants in common (D’aes et al., 2010). Biosurfactants are biologically produced amphiphilic molecules consisting of a hydrophilic head group and a hydrophobic moiety.

Leaf colonising Pseudomonads produce cyclic peptide biosurfactants (D’aes et al., 2010). Their ecophysiological role is not always clear, but it has been shown that Pseudomonads may gain different fitness advantages by producing surfactants including increasing survival during fluctuating humidity conditions on leaves (Burch et al., 2014) and by increasing local water availability due to the hygroscopic nature of their surfactants (Hernandez & Lindow, 2019). On agar plates it has been shown that biosurfactants increase surface mobility by swarming and it has been assumed that they may serve similar functions on leaves (Lindow & Brandl, 2003).

In this study, we characterised the physiological effect of biosurfactants in four different Pseudomonads that were isolated from leaves of spinach (*Pseudomonas* sp. FF1) or Romaine lettuce (*Pseudomonas* spp. FF2, FF3, and FF4) respectively. Their biosurfactants were characterised using mass spectrometry and their physical properties were analysed. Furthermore, we investigated the ecophysiological functions of the biosurfactants for the bacteria. To that end, random insertion libraries were produced and biosurfactant knockout mutants identified. The knockout mutants were characterised in a series of experiments that investigated fitness changes *in vitro* and *in planta*.

## Material and Methods

### Bacterial strains used in this study

Bacteria used in this study were *Pseudomonas* sp. FF1 (*P*FF1), *Pseudomonas* sp. FF2 (*P*FF2), *Pseudomonas* sp. FF3 (*P*FF3), *Pseudomonas* sp. FF4 (*P*FF4) (Burch et al., 2011); All Pseudomonads were kind gifts of Adrien Burch and Steven Lindow (UC Berkeley)) and *E. coli* Stellar (Lucigen). *P*FF1 was isolated from spinach, *P*FF2, *P*FF3, and *P*FF4 were isolated from Romaine lettuce. Pseudomonads were routinely grown on liquid King’s B (KB, 20 g proteose peptone, 1.15 g K_2_HPO_4_, 1.5 g Mg[SO_4_]*7H_2_O. 10 g glycerol per liter, pH 7; for agar medium KBA, add 15 g agar per liter) or Lysogeny Broth (LB, 5 g yeast extract, 10 g tryptone, 10 g NaCl per liter, pH 7; for agar medium add 15 g agar per liter). *E. coli* was routinely grown on LB and LBA. For *in planta* competition experiments, spontaneous streptomycin resistant mutants of the wildtype Pseudomonads were selected (Newcombe & Hawirko, 1949). Where appropriate, the media were supplemented with kanamycin (50 μg ml^-1^) or streptomycin (50 μg ml^-1^).

### 16S rRNA gene sequencing

To determine the phylogeny of the strains, their 16S rRNA gene was amplified from genomic DNA that was extracted using the NucleoSpin® Microbial DNA Kit (Macherey Nagel) following the manufacturer’s recommendations. A PCR using KAPA2G Fast 2x Ready Mix with Dye (Kapa) was performed using the manufacturer’s recommendation, 1 μL of genomic and 16S rRNA gene targeting primers SLK8-F 5’-AGAGTTTGATCATGGCTCAGAT-3’ and SRK1506-R 5’-TACCTTGTTACGACTTCACCCC-3’. Resulting ∼1.5 Kbp fragments were sequenced (Eurofins Genomic) and then curated and assembled using Geneious prime (Geneious). The assembled fragments were uploaded to ezbiocloud (Yoon et al., 2017) and the 30 best matches of organisms that were validly named were recovered for each of the four strains. Additional *Pseudomonas* 16S sequences and outgroup sequences were recovered from the silva database (Glöckner et al., 2017). All sequences were compiled into a fasta file and aligned and visualised using the FastME/OneClick option of ngphylogeny.fr (Lemoine et al., 2019). The resulting tree was imported into iTol, edited for publication and then exported (Letunic & Bork, 2019).

### Preparation of electrocompetent Pseudomonads

Electrocompetent Pseudomonads were produced as explained elsewhere (Artiguenave et al., 1997). Briefly, bacteria were grown overnight in 6 ml KB in a shaking incubator at 25 °C. Three ml of the overnight culture were then used to inoculate 100 ml KB that were incubated at 25 °C in a shaking incubator until the culture reached mid-exponential growth phase OD_600nm_ of approximately 0.6. The culture was then split in 50 ml aliquots and cooled on ice for 30 minutes. Bacteria were then harvested by centrifugation at 6000 *g* and 4 °C for 10 minutes. The supernatant was dismissed and the aliquots were washed twice with 50 ml ice-cold sterile water. Then they were washed in 25 ml ice-cold water and the aliquots were combined again. After a final centrifugation, the cell pellet was resuspended in 250 μl sterile 10% glycerol and distributed in 50 μl aliquots that were stored at -80 °C.

### Random transposon mutagenesis

Random knockout mutants were produced using the EZ::Tn5^Tm^ <KAN-2> Tnp Transposome^Tm^ kit (Epicentre) following the manufacturers recommendations. In brief, 50 μl electrocompetent Pseudomonads were thawed on ice and 1 μl Tn*5*-transposome and 1 μl endonuclease inhibitor were mixed with the cells. The mix was incubated for 5 minutes on ice before the cells were pipetted into a pre-chilled 0.1 cm gap electroporation cuvette. A gene pulser (Bio-Rad) was used to pulse the cells (2.5 kV, 200 Ω, 25 μF). Immediately after that, 1 ml SOC (SOB: 20 g tryptone, 5 g yeast extract, 0.5 g NaCl, 10 ml 250 mM KCl per liter, pH 7. SOC: SOB supplemented with 5 ml 2 M MgCl_2_ and 20 ml 1 M glucose) was added and the cells were incubated for 1 hour at 30 °C and 150 rpm. Transposon insertion mutants were selected on minimal medium agar plates (15 ml glycerol, 5 g L-glutamin, 1.5 g K_2_HPO_4_, 1.15 g MgSO_4_ × 7H_2_O, 15 g agar per liter, pH 7) supplemented with kanamycin. Minimal medium was used to prevent the growth of auxotrophic mutants. Transposon mutants could be detected after 2 days.

To determine the site of transposon integration, genomic DNA of knockout mutants was isolated using the ISOLATE II kit (Bioline). Genomic DNA was cut using KpnI (New England Biolabs) or EcoRI and ligated into similarly digested and dephosphorylated vector pUC19 (New England Biolabs) using T4-ligase (New England Biolabs) following the recommendations of the manufacturer. 5 μl per ligation mix were transformed into chemical competent *E. coli* Stellar using the manufacturers recommendations. Clones harboring plasmids containing the transposon were selected on LB supplemented with kanamycin. Inserts of the plasmids were sequenced using the transposon specific primer kan2_RP-1 (5’-gcaatgtaacatcagagattttgag-3’). Sequencing results were compared to the NCBI database using NCBI BLAST restricted to the genus Pseudomonas (Altschul et al., 1990).

### Screens for surfactant production

To screen for surfactant production, the atomised oil assay was performed (Burch et al., 2010). To that end, agar plates containing transposon mutants were sprayed with hydrophobic dodecan using an airbrush. Bacterial colonies that produced surfactants resulted in a halo around the colony where the surfactant in the agar changes the surface angle of oil droplets on the surface. Colonies that lacked this characteristic halo were further characterised. Presumptive surfactant mutants were tested in the drop collapse assay as described previously (Oso et al., 2019). Briefly, 2 μl of Magnatec 10W-40 oil (Castrol) were pipetted into each well of a 96-well plate lid (Corning incorporated) and were allowed to equilibrate for 2 hours to ensure that each well was evenly coated. Bacterial overnight cultures were centrifuged at 2600 × *g* for 10 minutes. Five μL of the culture supernatant was pipetted into the centre of an oil filled well. Drops that collapsed into the oil, i.e. decreased their contact angle, were positive for surfactant production while drops that remained intact and stayed on top of the oil were negative for surfactant production. All experiments were performed in at least 8 biological replicates.

### Extraction of surfactants

Bacterial strains were grown as crude streaks on five separate KBA plates for 48 hours at 25 °C. Afterwards, bacterial biomass was harvested using 5 ml of sterile water per plate and the cell suspensions of all 5 plates were combined in a 50 ml centrifugation tube. 25 ml ethyl acetate was added to the suspension and the tube was vortexed for 3 minutes. The mixture was then centrifuged for 10 minutes at 1000 x *g* to facilitate separation of the aqueous and organic phase. The organic phase was recovered using a glass pipette and transferred to a glass vessel before the ethyl acetate was evaporated off under constant nitrogen flow. The result was resolved in ethanol and sterile filtered through a 0.22 μm filter. The filtered solution was then dried under constant nitrogen flow and weight before it was resuspended to 5 μg ml^-1^ in ethyl acetate.

### Mass-spectrometric analysis

Mass spectrometric analysis of the biosurfactants was performed using a QTRAP 4500 (Applied Biosystems, AB Sciex) triple-quadrupole mass spectrometer, operated in negative electrospray ionization (ESI) – Q1 Scan Modus. The surfactant solution with a concentration of 5 μg ml^-1^ was injected via a syringe pump set to a flow rate of 10 μl min-1 directly into the MS. The analytes were detected in negative mode within a mass over charge range of 1000 – 1200 m/z.

### *Plant growth and* in planta *experiments*

*Arabidopsis thaliana* was grown axenically as described previously (Miebach et al., 2020). Briefly, Arabidopsis seeds were sterilised in a 1.5 ml Eppendorf tube by adding 1 mL 70 % ethanol and 0.1 % Triton X-100. The seeds were vortexed and then incubated for one minute. The supernatant was removed by pipetting, followed by the addition of 1 ml 10 % bleach and 10 μl of 0.1 % Triton X-100 for 12 minutes. After removing the bleach, the seeds were rinsed thrice with 1 ml of sterile distilled water were stratified for 48 hours at 4 °C. Stratified seeds were pipetted onto Murashige and Skoog-agar (MS-agar, 2.2 g of Murashige and Skoog medium including vitamins (Duchefa) and 10 g plant agar (Duchefa) per litre of milliQ water, pH 5.8) filled 200 μL pipette tips that were shortened by 1 cm to allow the plant’s roots to easily pass the tip. The tips were placed pointy end first into a MS-agar plate. The seeds were germinated for seven days at short day conditions (11 hours day/ 13 hours night). After the germination period, the seedling-filled tips were transferred to autoclaved Magenta™ GA-7 (bioWORLD) plant culture boxes filled with finely ground 90 g zeolite clay (Purrfit Clay Litter, Vitapet) and 60 ml MS medium. Four seedlings were transferred into each Magenta box and the plants were grown for an additional three weeks at short day conditions (11 hours day/ 13 hours night, chamber set to 85% relative humidity). To prepare bacterial inoculum, bacteria were cultured on LB broth overnight. Bacteria were then harvested by 10 min centrifugation at 2600 *g* and washed with 1 × phosphate buffer saline (PBS, 0.2 g L^−1^ NaCl, 1.44 g L^−1^ Na_2_ HPO_4_ and 0.24 g L^−1^ KH_2_ PO_4_). Bacteria were resuspended to an OD_600nm_ 0.5 and then serial diluted to OD_600nm_ 0.00005. For competition experiments wildtype and surfactant knockout strains were mixed at a ratio of 1:1. 100 μL of the mix or the monocultures were inoculated onto three week-old Arabidopsis using an T-180 airbrush (KKmoon).

Bacteria were recovered by harvesting the leaf material of individual plants, placing them in a 1.5 ml Eppendorf vial. The plants were weighed and 1 mL 1 × PBS were added. The vial was vortexed for 2 minutes and then sonicated for 5 minutes in a sonication bath (Elmasonic) before they were vortexed for another 2 minutes. The supernatant was serial diluted and CFU of wildtype and surfactant mutants were determined by growing the strains on LB agar containing appropriate antibiotics to select for either the spontaneous streptomycin resistant wildtype or the kanamycin resistant mutants.

### Diesel utilisation assay

To measure the ability of wildtype and surfactant knockout mutants to grow on diesel as the sole source of carbon, Bushnell-Haas broth (0.2 g L^−1^ MgSO_4_, 0.02 g L^−1^ CaCl_2_, 1.0 g L^−1^ KH_2_PO_4_, 1.0 g L^−1^ K_2_HPO_4_, 1.0 g L^−1^ NH_4_NO_3_ and 0.05 g L^−1^ FeCl_3_, pH 7.2), was supplemented with 1% diesel (commercial diesel, locally sourced) (Oso et al., 2019). Bushnell-Haas broth without additional carbon source was used as a negative control. In control experiments, to complement surfactant knockout mutants, between 0.23-0.265 mg mL^-1^ of isolated WT surfactants or 0.1 mg mL^-1^ Tween-20 were supplemented. Bacteria were grown overnight in LB, diluted 100 × using Bushnell-Haas broth without carbon source. The diluted bacterial suspensions were inoculated into 50 mL broth cultures in 250 mL Erlenmeyer flasks. Cultures were incubated at 30°C and 200 rounds per minutes for up to 17 days. Cell density was regularly measured by determining the optical density at 600 nm using a spectrophotometer (Biochrom WPA CO8000, Biowave). All experiments were performed in three biological replicates.

## Results

### *Phylogenetic placement of* Pseudomonas *sp. FF1, FF2, FF3 and FF4*

Analysis of the 16S rRNA genes of all four isolates revealed that they are all members of the genus Pseudomonas and members of the *Pseudomonas fluorescens* lineage and subgroup (Peix et al., 2018). *P*FF1 clusters closely with *Pseudomonas orientalis, P*FF2 clusters closely with *Pseudomonas extremaustralis*, while *P*FF3 and *P*FF4 cluster closely with *Pseudomonas paralactis* (Figure 1). *P*FF1 and *P*FF2 are closer related to each other than to *P*FF3 and *P*FF4. *P*FF3 and *P*FF4 are closely related.

**Figure 1.**
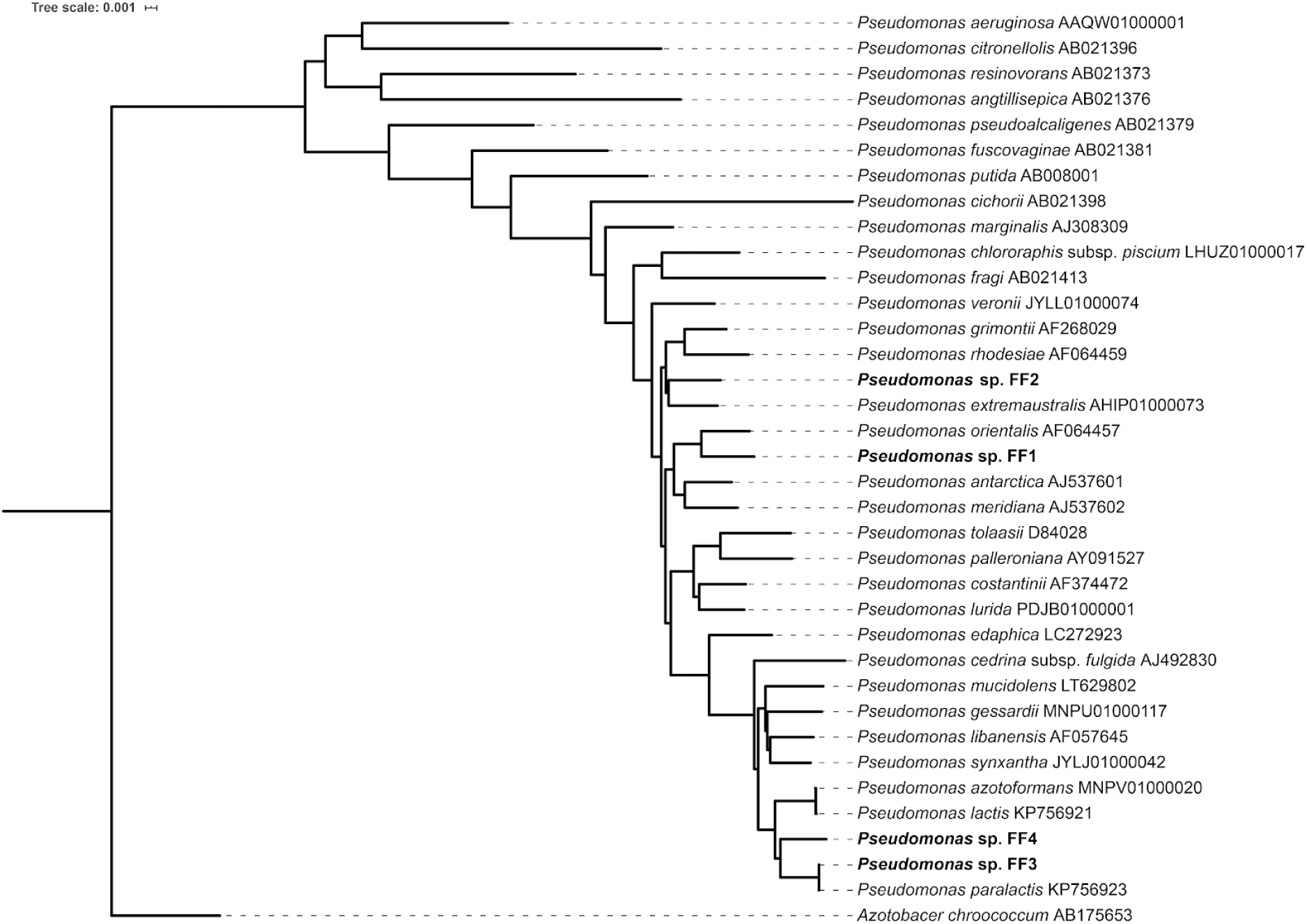
Phylogenetic placement of the four isolated Pseudomonads. The newly sequenced isolates are highlighted in bold. NCBI accession numbers of the respective sequences are noted behind the species names. *Azotobacter chroococcum* was used as an outgroup.

### Surfactant production of tested Pseudomonads

All four wildtype Pseudomonads were tested for their production of surfactants on agar plates using the atomised oil assay (Burch et al., 2010; Oso et al., 2019). All four strains produced clear halos where the reflection of the oil to light changed indicating production of surfactants (Figure 2A-D). Similarly, the positive control Tween-20 showed a halo (Figure 2E), while the negative control, *E. coli* Dh5α, was lacking a halo (Figure 2F). The drop collapse assay was used as a secondary test for surfactant production. All tested wildtype culture supernatants collapsed into the engine oil (Figure 2G-J). The collapse is due to a change in surface tension of the supernatant.

**Figure 2.**
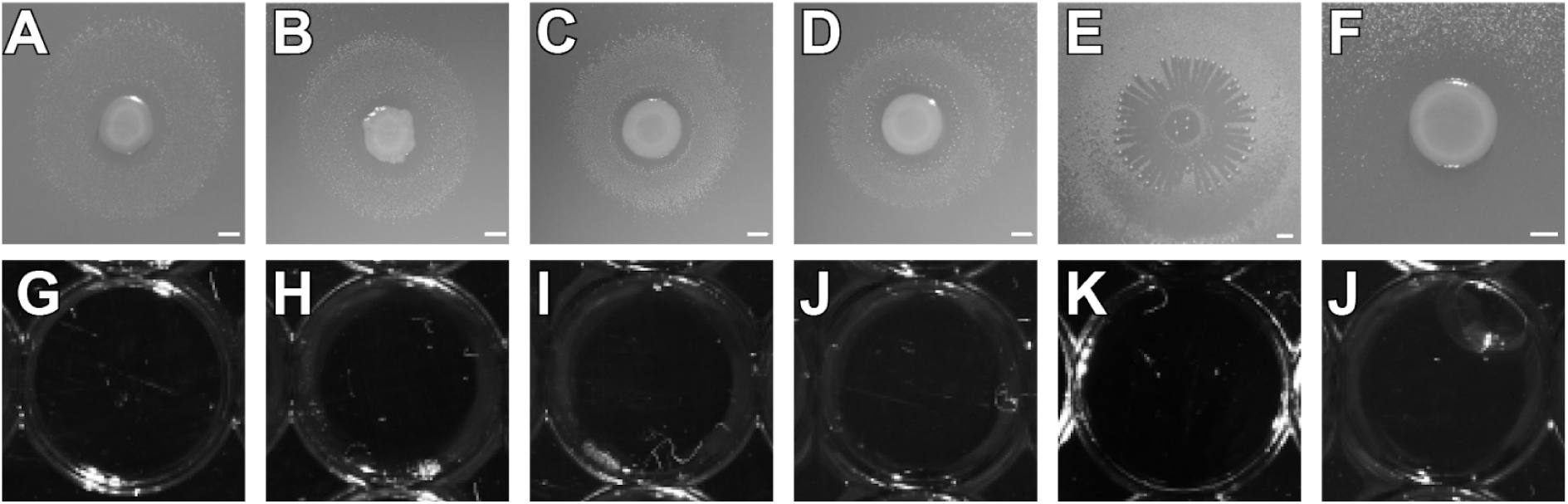
A-F) Atomised oil assays to demonstrate the production of surfactants. **A-D)** wildtype colonies of *P*FF1, *P*FF2, *P*FF3, and *P*FF4, respectively, exhibiting a halo indicative for surfactant production. **E)** Tween-20 **F)** E. coli Dh5α **G-L) Drop collapse assays to demonstrate the production of surfactants**. Culture supernatants of wildtype *P*FF1, *P*FF2, *P*FF3, and *P*FF4, respectively, collapsed into oil indicative for surfactant production. **K)** collapsed drop containing Tween-20. **L)** Non-collapsed drop if *E. coli* culture supernatant.

### Mass spectrometric analysis surfactants

The analysis of surfactants harvested from the Pseudomonads using LC-MS with ESI in negative mode revealed that *P*FF1 and *P*FF2 produced the same compounds with a characteristic main peak at m/z=1124.59, which can be attributed to the deprotonated molecular ion [M-H]^-^. The analogous pattern for the protonated molecular ion [M+H]^+^ has been previously described for the cyclic lipopeptide viscosin when using ESI in positive mode for detection(de Bruijn et al., 2008; Laycock et al., 1991). Similarly, *P*FF3 and *P*FF4 share the same characteristic main peak at m/z=1138.60, the analogous pattern has previously been described for the cyclic lipopeptide massetolide A(de Bruijn et al., 2008).

### Random Tn*5* mutagenesis and mutant characterisation

The surfactants producing wildtypes were subjected to random insertion mutagenesis using the EZ-Tn5 transposon system. The screen resulted in a transposon mutant library with several hundred transposon mutants for each of the four isolates. Each of the mutant libraries was screened with the atomised oil assay for lack of surfactant production mutants. Mutants lacking surfactant production were identified and one mutant for each isolate was selected for further characterisation (Figure 4A-D). The drop collapse assay was conducted and confirmed the results of the atomised oil assay (Figure 4E-H). The insertion site of each mutant was determined by digesting the genomic DNA of the mutants and cloning it into pUC19 before selecting for kanamycin resistance encoded in the transposon (Supplemental Table 1).

**Figure 3.**
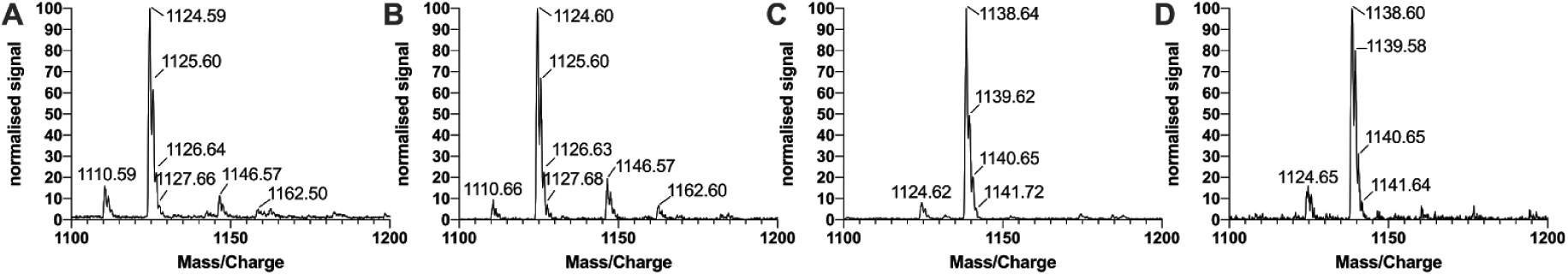
A-D) MS/MS spectra of extracted surfactants of. *P*FF1, *P*FF2, *P*FF3, and *P*FF4 respectively. *P*FF1 and *P*FF2 both produce viscosin, *P*FF3 and *P*FF4 both produce massetolide A. Spectra were normalised against the maximal intensity.

**Figure 4.**
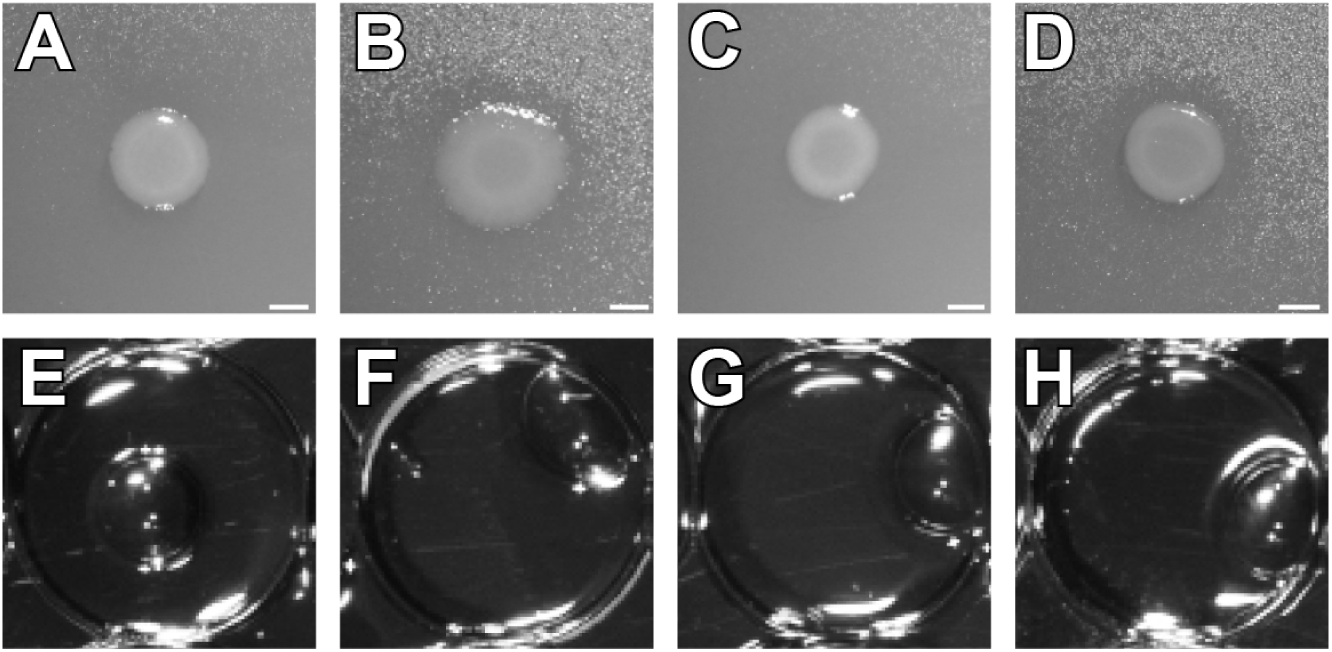
A-D) Atomised oil assay to demonstrate the production of surfactants. Tn5-transposon insertion mutant colonies *P*FF1::ezTn5-visB, *P*FF2::ezTn5-visB, *P*FF3::ezTn5-massB, and *P*FF4::ezTn5-massB, respectively, lacking a halo indicative for surfactant production. **E-H) drop collapse assays to demonstrate the production of surfactants**. Culture supernatants of Tn5-transposon insertion mutant *P*FF1::ezTn5-visB, *P*FF2::ezTn5-visB, *P*FF3::ezTn5-massB, and *P*FF4::ezTn5-massB, respectively, showing a beaded bubble swimming on top of oil, indicative for the lack of surfactants.

The investigated *P*FF1 mutant carried an insertion in a gene with 97% similarity to a non-ribosomal peptide synthetase in *P. orientalis* F9 (Genbank: BOP93_14875) (Zengerer et al., 2018) which has an 80% peptide similarity to the *viscB* gene of *P. fluorescens* SBW25 (UniProtKB ID: C3K9G2) (De Bruijn et al., 2007; Silby et al., 2009). The investigated *P*FF2 mutant carried an insertion in a gene with a 86% similarity to the *viscB* gene (Genbank: CAY48788.1) of *P. fluorescens* SBW25, respectively. Therefore, they are designated *P*FF1::ezTn5-viscB and *P*FF2::ezTn5-viscB, respectively. The *viscB* gene encodes for a non-ribosomal peptide synthetase that, in conjunction with *viscA* and *viscC*, produces the cyclic lipopeptide biosurfactant viscosin (De Bruijn et al., 2007). The *P*FF3 Tn*5* transposon mutant carried an insertion in a gene with 99% similarity to the *massB* gene in *Pseudomonas fluorescens* SS101 (Genbank: ABH06368.2). The *P*FF4 Tn*5* transposon mutant carried an insertion in a gene with 95% similarity to the *massB* gene in *Pseudomonas fluorescens* SS101 (de Bruijn et al., 2008). The *massB* gene is part of the massetolide A synthesis gene cluster. Therefore the mutants were designated *P*FF3::ezTn5-massB and *P*FF4::ezTn5-massB.

After extracting agar plates to recover surfactants for the analysis using mass spectrometry, no surfactants could be detected (Figure 5 A-D).

**Figure 5.**
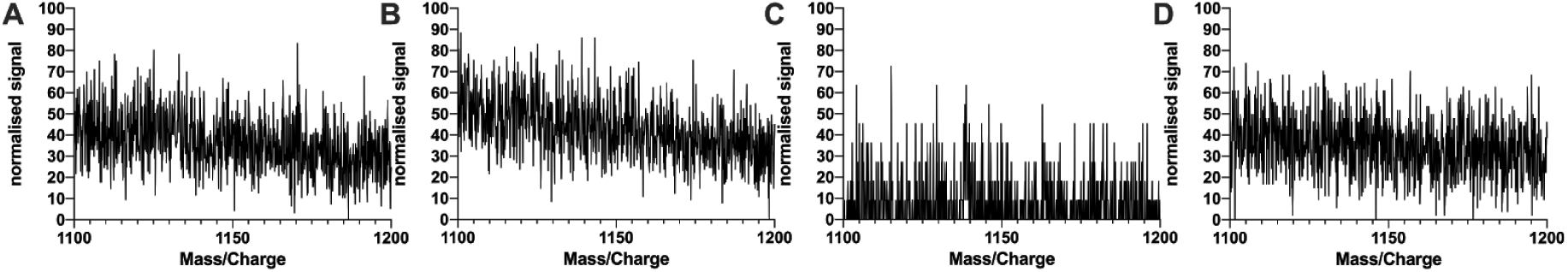
A-D) Knockout mutants show no sign of surfactant production - MS/MS spectra of extracts of *P*FF1::ezTn5-viscB, *P*FF2::ezTn5-viscB, *P*FF3::ezTn5-massB, and *P*FF4::ezTn5-massB, respectively. None of the random knockout mutants produced detectable surfactant peaks at the respective wildtype m/z values. Spectra were normalised against the maximal intensity.

The effect of the transposon insertions and the lack of surfactant production was tested in shaking liquid cultures in two different conditions, either KB complex medium (Supplemental Figure 1A), or M9 minimal medium supplemented with glucose as the sole source of carbon (Supplemental Figure 1B). None of the tested insertion mutants exhibited significantly changed doubling times under the two tested conditions.

### Growth on diesel oil as sole carbon source

To investigate if the lack of surfactant production could impact the ability of the Pseudomonad strains to degrade alkanes, the different wildtypes and transposon mutants were grown on Bushnell-Haas broth with diesel as the sole carbon source. This experiment revealed that all surfactant mutants, even though they were still able to grow on diesel, had a reduced growth rate, and a reduced final optical density after up to 21 days of growth (Figure 6). No growth could be observed on Bushnell-Haas broth without carbon source for either the wildtype or the surfactant mutants (data not shown). In general, the growth on diesel oil was slower compared to growth on complex medium or minimal medium supplemented with glucose as sole carbon source and better described by a linear function than an exponential growth function. By supplementing knockout mutants growing on diesel with biosurfactants harvested from respective wildtype strains or Tween-20 growth on diesel could be complemented in parts or completely compared to the wildtype. Knockout mutants could not grow on provided surfactants to a degree that explains the increased growth on diesel (Supplemental figure 2).

**Figure 6.**
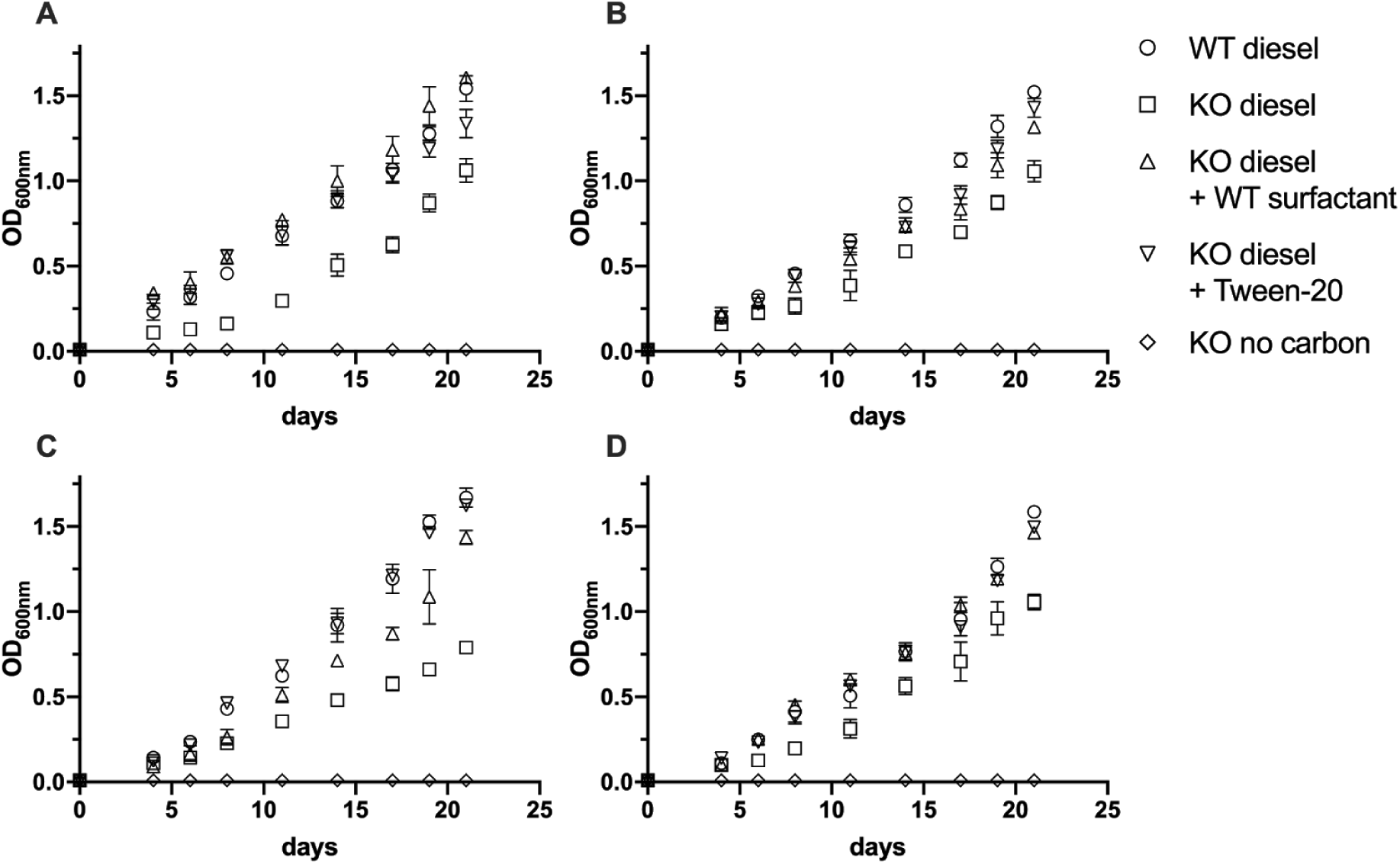
Utilisation of diesel by biosurfactant knockout mutants and wildtypes. A) *P*FF1, B) *P*FF2, C) *P*FF3, D) *P*FF4. Each wildtype and knockout mutant was grown in Bushnell-Haas broth supplemented with diesel as the sole source of carbon (circle and square, respectively). Knockout mutants were complemented with either wildtype surfactant (triangle), Tween-20 (inverted triangle) or were incubated with no additional carbon source (diamond). Error bars depict the standard deviation of the mean.

### Fitness *in planta*

To investigate changes in the ability of the transposon mutants to colonise leaf surfaces, the mutants were co-inoculated with the respective wildtypes by airbrushing. Whole above-ground plant material was sampled daily for six days and colony forming units of wildtype and transposon mutants were determined (Figure 7). The initial bacterial densities were similar between wildtype and knockout mutants. wildtypes (PFF1, PPF2, PPF3 and PPF4) and corresponding mutants (PFF1::ezTn5-viscB, PFF2::ezTn5-viscB, PFF3::ezTn5-massB and PFF4::ezTn5-massB) colonised Arabidopsis at similar rates. PPF1, PPF2 and their mutants reached approximately 10^7^ CFU per gram of plant weight, whereas PFF3, PPF4 and their mutants reached approximately 10^6^ CFU per gram of plant weight. Thus, no differences between the plant colonisation of wildtype and mutants were found. Furthermore, growth in planta of all strains was tested individually, no significant differences in plant colonisation could be determined (Supplemental figure 3).

**Figure 7.**
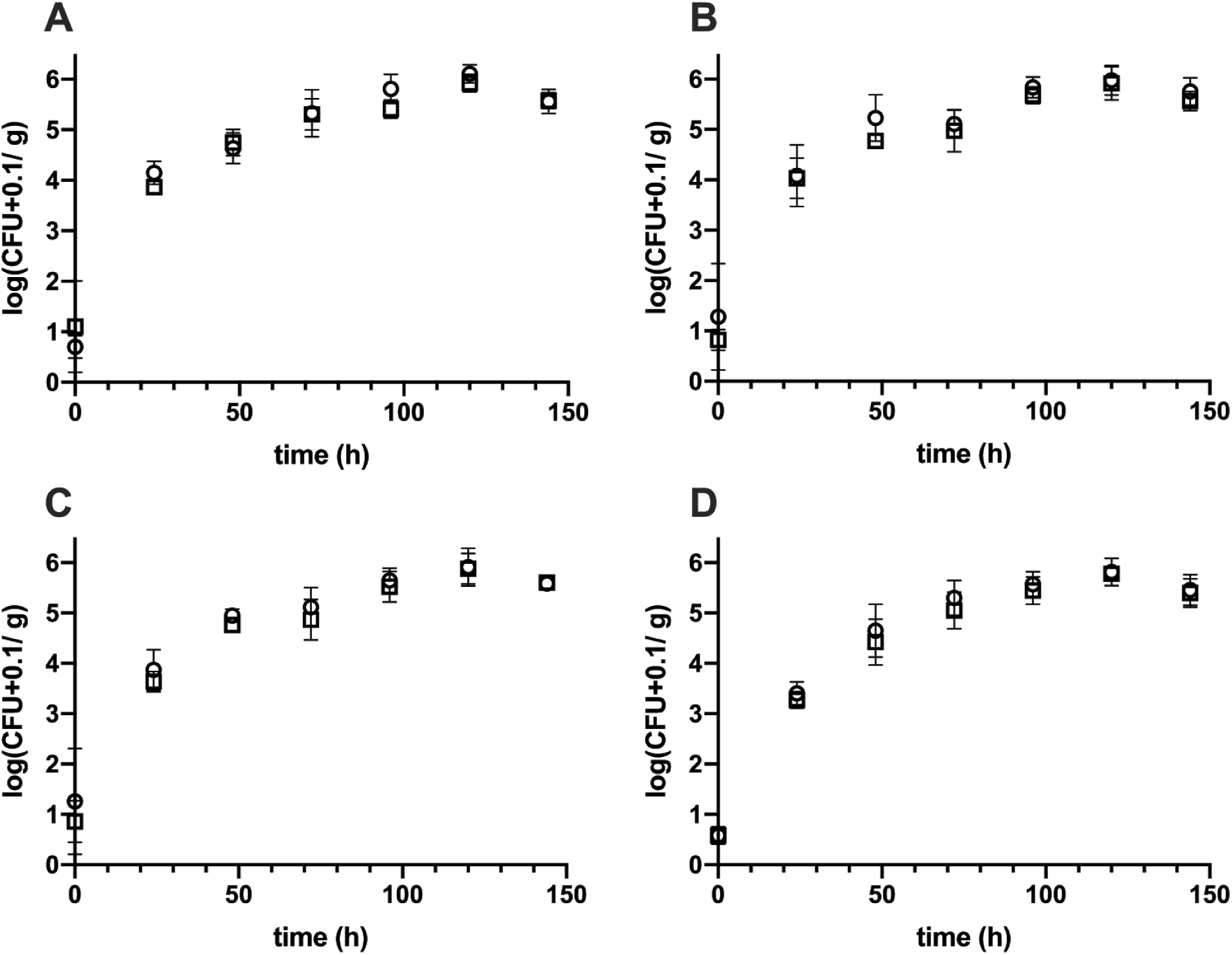
*In planta* competition of wildtypes (open circles) and mutants (open squares). A) *P*FF1 vs. *P*FF1::ezTn5-visB, B) *P*FF2 vs. *P*FF2::ezTn5-visB, C) *P*FF3 vs. *P*FF3::ezTn5-massB, D) *P*FF4 vs. *P*FF2::ezTn5-massB. wildtypes are depicted by circles, knockout mutants by squares. Error bars depict the standard deviation of the mean.

## Discussion

All four Pseudomonads isolated from either spinach or romaine lettuce leaf material (Burch et al., 2016) belong to the fluorescent Pseudomonads (Gomila et al., 2015). *P*FF1 and *P*FF2 are phylogenetically more closely related to each other than to *P*FF3 and *P*FF4. *P*FF3 and *P*FF4 are very closely related. All four strains are produced surfactants on agar plates and in liquid culture as shown by the atomised oil assay and the drop collapse assay. As the ability to produce surfactants is widely distributed in the genus *Pseudomonas*, this result was not surprising (Geudens & Martins, 2018; Nybroe & Sørensen, 2004). The relatedness of the strains is also reflected in the surfactants that each of the strains is producing: *P*FF1 and *P*FF2 produce the viscosin-like surfactants, while *P*FF3 and *P*FF4 are produced massetolide A-like surfactants. The production of viscosin and massetolide A by Pseudomonads has been demonstrated previously (de Bruijn et al., 2008). Both viscosin and massetolide A are the product of nonribosomal peptide synthetase genes. Viscosin production depends on a gene cluster encompassing the three genes *viscA, viscB, and viscC* and which spans approximately 32 kb (De Bruijn et al., 2007). Massetolide A production also depends on a gene cluster which encompasses the three genes *massA, massB* and *massC* and spans approximately 30 kb (de Bruijn et al., 2008).

To further investigate the ecological function of the surfactants in the leaf colonising Pseudomonads, random Tn*5* transposon insertion mutants were produced and further characterised. The screen yielded complete loss of surfactant production mutants for every strain, indicating that each strain only encodes for one surfactant that is active during the selection conditions. The insertion sites were mapped to genes that matched previously characterised non-ribosomal peptide synthase clusters responsible for surfactant production, and which matched the surfactants that were identified using mass-spectrometry. *P*FF1 and *P*FF2 knockout mutants were mapped to *viscB* gene homologues, and *P*FF3 and *P*FF4 knockout to *massB* gene homologues (De Bruijn et al., 2007; de Bruijn et al., 2008).

The assumption that only one surfactant is produced by each strain was corroborated by a sequence of experiments during which the surfactant mutants consistently failed to produce signs of surfactant production independent of their growth conditions. The surfactant mutants failed to produce halos in the atomised oil assay, and the culture supernatant did not collapse into motor oil in the drop collapse assay. Mass spectrometric analysis of the knockout mutants showed that the production of surfactants was completely abolished and no detectable peak pattern was found after the surfactant extraction procedure (Figure 5).

Despite the loss of surfactant production and the additional burden of expressing the kanamycin resistance gene from the Tn*5* transposon, the insertions had no detectable fitness effects in either complex KB medium or minimal M9 medium supplemented with glucose. In shaking liquid cultures, surfactants did not provide critical functions for growth (Supplemental figure 1). We hypothesise that surfactants may enable bacteria to utilise parts of the plant cuticle as a source for carbon. Even though it was not possible to show that Pseudomonads and their respective mutants had differential abilities to utilise hydrocarbon components from isolated cuticles (data not shown), a clear difference in the ability of wildtype and mutant to utilise diesel for growth was demonstrated (Figure 6). Even though growth was not completely abolished, it was markedly reduced. This could also explain why growth on isolated cuticles did not yield conclusive results and differences between wildtype and knockout mutant. Due to the size of the non-ribosomal peptide synthetase genes, it was not possible to construct rescue mutants. However, we attempted to complement the reduced ability of the knockout mutants to degrade diesel oil by adding harvested wildtype surfactant or Tween-20 to growing cultures. Indeed, both surfactants were able to complement the growth phenotype either in parts or completely, evidencing that the lack of surfactants was the causal reason for reduced growth. Despite the chain length differences between the diesel (Wante & Leung, 2018) and the alkane monomers in waxes of leaf cuticles (Zeisler-Diehl et al., 2018), the chemistry of both aliphatic mixtures contain similar monomers. It is thus not unthinkable that, under nutrient limiting conditions, the *Pseudomonas* strains tested here are able to utilise aliphatic components of leaf cuticles in a surfactant-dependent manner. However, we failed to provide a final proof of this relationship. To investigate the role of the surfactants during plant colonisation we inoculated axenically grown Arabidopsis with mixtures of wildtype and knockout mutants or with individual strains. During co-inoculation with their respective wildtypes onto axenic Arabidopsis, no fitness disadvantages for the knockout mutants were detected. This might be a consequence of the surfactant acting as a public good that increases the fitness of wildtype and co-inoculated mutants alike (Lyons & Kolter, 2017). However, single strain inoculations also did not result in a diminished ability of the knockout mutants to colonise Arabidopsis. This is in contrast to previous experiments that demonstrated that surfactants do indeed have a positive effect on plant colonisation (Burch et al., 2014). It is noteworthy that the experimental setup used in our study was markedly different including a different plant host as well as incubation conditions under constant relative humidities. While previously it was shown that fluctuating humidities are a prerequisite to result in a fitness advantage. Therefore, it might still be possible the surfactants in the here-tested strains will impact plant colonisation for example under fluctuating relative humidities.

## Conclusion

The experiments reported here demonstrated that the biosurfactants produced by four different leaf colonising Pseudomonads impacted on their ability to degrade aliphatic compounds. However, the ability to produce biosurfactants had no measurable impact on the ability of the strains to colonise axenic Arabidopsis leaves in competition or after individual strain inoculations. We gathered additional evidence that the bacteria may utilise aliphatic compounds originating from leaf cuticles but failed to conclusively demonstrate a relationship between surfactant production and leaf colonisation ability. Future studies will have to be performed to address this hypothesis.

## Acknowledgements

We thank Adrien Burch and Steven Lindow, UC Berkeley, for the kind gift of the *Pseudomonas* strains. We thank Paula Jameson for critically reading the manuscript and Thomas Evans for technical support.

## References

Altschul, S. F., Gish, W., Miller, W., Myers, E. W., & Lipman, D. J. (1990). Basic local alignment search tool. Journal of Molecular Biology, 215(3), 403–410.

Artiguenave, F., Vilaginès, R., & Danglot, C. (1997). High-efficiency transposon mutagenesis by electroporation of a Pseudomonas fluorescens strain. FEMS Microbiology Letters, 153(2), 363–369.

Burch, A. Y., Browne, P. J., Dunlap, C. A., Price, N. P., & Lindow, S. E. (2011). Comparison of biosurfactant detection methods reveals hydrophobic surfactants and contact-regulated production. Environmental Microbiology, 13(10), 2681–2691.

Burch, A. Y., Do, P. T., Sbodio, A., Suslow, T. V., & Lindow, S. E. (2016). High-level culturability of epiphytic bacteria and frequency of biosurfactant producers on leaves. Applied and Environmental Microbiology, 82(19), 5997–6009.

Burch, A. Y., Shimada, B. K., Browne, P. J., & Lindow, S. E. (2010). Novel high-throughput detection method to assess bacterial surfactant production. Applied and Environmental Microbiology, 76(16), 5363–5372.

Burch, A. Y., Zeisler, V., Yokota, K., Schreiber, L., & Lindow, S. E. (2014). The hygroscopic biosurfactant syringafactin produced by Pseudomonas syringae enhances fitness on leaf surfaces during fluctuating humidity. Environmental Microbiology, 16(7), 2086–2098.

Buschhaus, C., & Jetter, R. (2011). Composition differences between epicuticular and intracuticular wax substructures: How do plants seal their epidermal surfaces? Journal of Experimental Botany, 62(3), 841–853.

Cabrefiga, J., Bonaterra, A., & Montesinos, E. (2007). Mechanisms of antagonism of Pseudomonas fluorescens EPS62e against Erwinia amylovora, the causal agent of fire blight. International Microbiology: The Official Journal of the Spanish Society for Microbiology, 10(2), 123–132.

D’aes, J., De Maeyer, K., Pauwelyn, E., & Höfte, M. (2010). Biosurfactants in plant-Pseudomonas interactions and their importance to biocontrol. Environmental Microbiology Reports, 2(3), 359–372.

De Bruijn, I., de Kock, M. J. D., de Waard, P., van Beek, T. A., & Raaijmakers, J. M. (2008). Massetolide A biosynthesis in Pseudomonas fluorescens. Journal of Bacteriology, 190(8), 2777–2789.

De Bruijn, I., de Kock, M. J. D., Yang, M., de Waard, P., van Beek, T. A., & Raaijmakers, J. M. (2007). Genome-based discovery, structure prediction and functional analysis of cyclic lipopeptide antibiotics in Pseudomonas species. Molecular Microbiology, 63(2), 417–428.

Freimoser, F. M., Pelludat, C., & Remus-Emsermann, M. N. P. (2016). Tritagonist as a new term for uncharacterised microorganisms in environmental systems. The ISME Journal, 10(1), 1–3.

Geudens, N., & Martins, J. C. (2018). Cyclic lipodepsipeptides from Pseudomonas spp. - Biological Swiss-army knives. Frontiers in Microbiology, 9, 1867.

Glöckner, F. O., Yilmaz, P., Quast, C., Gerken, J., Beccati, A., Ciuprina, A., Bruns, G., Yarza, P., Peplies, J., Westram, R., & Ludwig, W. (2017). 25 years of serving the community with ribosomal RNA gene reference databases and tools. Journal of Biotechnology, 261, 169–176.

Gomila, M., Peña, A., Mulet, M., Lalucat, J., & García-Valdés, E. (2015). Phylogenomics and systematics in Pseudomonas. Frontiers in Microbiology, 6, 214.

Graça, J. (2002). Glycerol and glyceryl esters of ω-hydroxyacids in cutins. Phytochemistry, 61(2), 205–215.

Hernandez, M. N., & Lindow, S. E. (2019). Pseudomonas syringae increases water availability in leaf microenvironments via production of hygroscopic syringafactin. Applied and Environmental Microbiology, 85(18). https://doi.org/10.1128/AEM.01014-19

Jeffree, C. E. (2006). The Fine Structure of the Plant Cuticle. In M. Riederer & C. Müller (Eds.), Biology of the Plant Cuticle (pp. 11–125). Blackwell Publishing Ltd.

Jetter, R., Kunst, L., & Samuels, A. L. (2006). Composition of Plant Cuticular Waxes. In M. Riederer & C. Müller (Eds.), Biology of the Plant Cuticle (pp. 145–181). Blackwell Publishing Ltd.

Kertesz, M. A., & Kawasaki, A. (2010). Hydrocarbon-Degrading Sphingomonads: Sphingomonas, Sphingobium, Novosphingobium, and Sphingopyxis. In K. N. Timmis (Ed.), Handbook of Hydrocarbon and Lipid Microbiology (pp. 1693–1705). Springer Berlin Heidelberg.

Kolattukudy, P. E. (1980). Biopolyester membranes of plants: cutin and suberin. Science, 208(4447), 990–1000.

Laycock, M. V., Hildebrand, P. D., Thibault, P., Walter, J. A., & Wright, J. L. C. (1991). Viscosin, a potent peptidolipid biosurfactant and phytopathogenic mediator produced by a pectolytic strain of Pseudomonas fluorescens. Journal of Agricultural and Food Chemistry, 39(3), 483–489.

Lemoine, F., Correia, D., Lefort, V., Doppelt-Azeroual, O., Mareuil, F., Cohen-Boulakia, S., & Gascuel, O. (2019). NGPhylogeny.fr: new generation phylogenetic services for non-specialists. Nucleic Acids Research, 47(W1), W260–W265.

Letunic, I., & Bork, P. (2019). Interactive Tree Of Life (iTOL) v4: recent updates and new developments. Nucleic Acids Research, 47(W1), W256–W259.

Lindow, S. E., & Brandl, M. T. (2003). Microbiology of the phyllosphere. Applied and Environmental Microbiology, 69(4), 1875–1883.

Lyons, N. A., & Kolter, R. (2017). Bacillus subtilis protects public goods by extending kin discrimination to closely related species. mBio, 8(4). https://doi.org/10.1128/mBio.00723-17

Miebach, M., Schlechter, R. O., Clemens, J., Jameson, P. E., & Remus-Emsermann, M. N.P. (2020). Litterbox-A gnotobiotic Zeolite-Clay System to Investigate Arabidopsis-Microbe Interactions. Microorganisms, 8(4), 464.

Newcombe, H. B., & Hawirko, R. (1949). Spontaneous mutation to streptomycin resistance and dependence in Escherichia coli. Journal of Bacteriology, 57(5), 565–572.

Nybroe, O., & Sørensen, J. (2004). Production of cyclic lipopeptides by fluorescent Pseudomonads. In J.-L. Ramos (Ed.), Pseudomonas: Volume 3 Biosynthesis of Macromolecules and Molecular Metabolism (pp. 147–172). Springer US.

Oso, S., Walters, M., Schlechter, R. O., & Remus-Emsermann, M. N. P. (2019). Utilisation of hydrocarbons and production of surfactants by bacteria isolated from plant leaf surfaces. FEMS Microbiology Letters, 366(6), fnz061.

Peix, A., Ramírez-Bahena, M.-H., & Velázquez, E. (2018). The current status on the taxonomy of Pseudomonas revisited: An update. Infection, Genetics and Evolution: Journal of Molecular Epidemiology and Evolutionary Genetics in Infectious Diseases, 57, 106–116.

Pizzolante, G., Durante, M., Rizzo, D., Di Salvo, M., Tredici, S. M., Tufariello, M., De Paolis, A., Talà, A., Mita, G., Alifano, P., & De Benedetto, G. E. (2018). Characterization of two Pantoea strains isolated from extra-virgin olive oil. AMB Express, 8(1), 113.

Pollard, M., Beisson, F., Li, Y., & Ohlrogge, J. B. (2008). Building lipid barriers: biosynthesis of cutin and suberin. Trends in Plant Science, 13(5), 236–246.

Remus-Emsermann, M. N. P., Lücker, S., Müller, D. B., Potthoff, E., Daims, H., & Vorholt, J. A. (2014). Spatial distribution analyses of natural phyllosphere-colonizing bacteria on Arabidopsis thaliana revealed by fluorescence in situ hybridization. Environmental Microbiology, 16(7), 2329–2340.

Remus-Emsermann, M. N. P., Schmid, M., Gekenidis, M.-T., Pelludat, C., Frey, J. E., Ahrens, C. H., & Drissner, D. (2016). Complete genome sequence of Pseudomonas citronellolis P3B5, a candidate for microbial phyllo-remediation of hydrocarbon-contaminated sites.Standards in Genomic Sciences, 11, 75.

Riederer, M., & Schreiber, L. (2001). Protecting against water loss: analysis of the barrier properties of plant cuticles. Journal of Experimental Botany, 52(363), 2023–2032.

Salam, L. B., Obayori, O. S., & Raji, S. A. (2015). Biodegradation of used engine oil by a methylotrophic bacterium, Methylobacterium mesophilicum isolated from tropical hydrocarbon-contaminated soil. Petroleum Science and Technology, 33(2), 186–195.

Samuels, L., Kunst, L., & Jetter, R. (2008). Sealing plant surfaces: Cuticular wax formation by epidermal cells. Annual Review of Plant Biology, 59(1), 683–707.

Schlechter, R. O., Miebach, M., & Remus-Emsermann, M. N. P. (2019). Driving factors of epiphytic bacterial communities: A review. Journal of Advertising Research, 19, 57–65.

Schmid, M., Frei, D., Patrignani, A., Schlapbach, R., Frey, J. E., Remus-Emsermann, M. N. P., & Ahrens, C. H. (2018). Pushing the limits of de novo genome assembly for complex prokaryotic genomes harboring very long, near identical repeats. Nucleic Acids Research, 46(17), 8953–8965.

Serrano, M., Coluccia, F., Torres, M., L’Haridon, F., & Métraux, J.-P. (2014). The cuticle and plant defense to pathogens. Frontiers in Plant Science, 5. https://doi.org/10.3389/fpls.2014.00274

Shepherd, T., & Wynne Griffiths, D. (2006). The effects of stress on plant cuticular waxes. The New Phytologist, 171(3), 469–499.

Silby, M. W., Cerdeño-Tárraga, A. M., Vernikos, G. S., Giddens, S. R., Jackson, R. W., Preston, G. M., Zhang, X.-X., Moon, C. D., Gehrig, S. M., Godfrey, S. A. C., Knight, C. G., Malone, J. G., Robinson, Z., Spiers, A. J., Harris, S., Challis, G. L., Yaxley, A. M., Harris, D., Seeger, K., … Thomson, N. R. (2009). Genomic and genetic analyses of diversity and plant interactions of Pseudomonas fluorescens. Genome Biology, 10(5), R51.

Wante, S. P., & Leung, D. W. M. (2018). Phytotoxicity testing of diesel-contaminated water using Petunia grandiflora Juss. Mix F1 and Marigold-Nemo Mix (Tagetes patula L.). Environmental Monitoring and Assessment, 190(7), 408.

Wattendorff, J., & Holloway, P. J. (1980). Studies on the Ultrastructure and Histochemistry of Plant Cuticles: The Cuticular Membrane of Agave americana L. in situ. Annals of Botany, 46(1), 13–28.

Xin, X.-F., Kvitko, B., & He, S. Y. (2018). Pseudomonas syringae: what it takes to be a pathogen. Nature Reviews. Microbiology, 16(5), 316–328.

Yeats, T. H., Buda, G. J., Wang, Z., Chehanovsky, N., Moyle, L. C., Jetter, R., Schaffer, A. A., & Rose, J. K.C. (2012). The fruit cuticles of wild tomato species exhibit architectural and chemical diversity, providing a new model for studying the evolution of cuticle function. The Plant Journal, 69(4), 655–666.

Yoon, S.-H., Ha, S.-M., Kwon, S., Lim, J., Kim, Y., Seo, H., & Chun, J. (2017). Introducing EzBioCloud: a taxonomically united database of 16S rRNA gene sequences and whole-genome assemblies. International Journal of Systematic and Evolutionary Microbiology, 67(5), 1613–1617.

Zeisler-Diehl, V. V., Barthlott, W., & Schreiber, L. (2018). Plant Cuticular Waxes: Composition, Function, and Interactions with Microorganisms. In H. Wilkes (Ed.), Hydrocarbons, Oils and Lipids: Diversity, Origin, Chemistry and Fate (pp. 1–16). Springer International Publishing.

Zengerer, V., Schmid, M., Bieri, M., Müller, D. C., Remus-Emsermann, M. N. P., Ahrens, C. H., & Pelludat, C. (2018). Pseudomonas orientalis F9: A potent antagonist against phytopathogens with phytotoxic effect in the apple flower. Frontiers in Microbiology, 9, 145.

